# Mass Spectrometry Imaging of N-Glycans Reveals Racial Discrepancies in Low Grade Prostate Tumors

**DOI:** 10.1101/2020.08.20.260026

**Authors:** Lindsey R. Conroy, Lyndsay E.A. Young, Alexandra E. Stanback, Grant L. Austin, Jinpeng Liu, Jinze Liu, Derek B. Allison, Ramon C. Sun

## Abstract

Prostate cancer is the most common cancer in men worldwide. Despite its prevalence, there is a critical knowledge gap regarding the underlining molecular events that result in higher incidence and mortality rate in Black men. Identifying molecular features that separate racial disparities is a critical step in prostate cancer research that could lead to predictive biomarkers and personalized therapy. N-linked glycosylation is a co-translational event during protein folding that modulates a myriad of cellular processes. Recently, aberrant N-linked glycosylation has been reported in prostate cancers. However, the full clinical implications of dysregulated glycosylation in prostate cancer has yet to be explored. Herein, we performed high-throughput matrix-assisted laser desorption ionization mass spectrometry analysis to characterize the N-glycan profile from tissue microarrays of over 100 patient tumors with over 10 years of follow up data. We identified several species of N-glycans that were profoundly different between low grade prostate tumors resected from White and Black patients. Further, these glycans predict opposing overall survival between White and Black patients with prostate cancer. These data suggest differential N-linked glycosylation underline the racial disparity of prostate cancer prognosis. Our study highlights the potential applications of MALDI-MSI for digital pathology and biomarker to study racial disparity of prostate cancer patients.

**Graphical Abstract:** 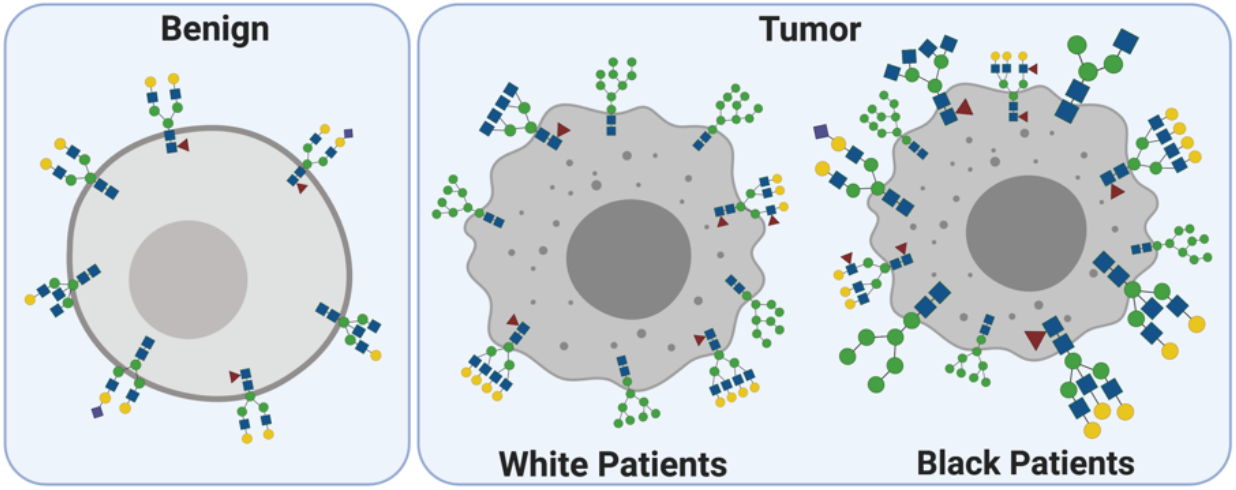

## Introduction

Prostate cancer is the most common cancer in men and is the second leading cause of cancer-related mortality in men worldwide (1). Many factors contribute to the development and progression of prostate cancer including age, family history, ethnicity, and diet or lifestyle (2, 3). Patient prognosis largely depends on tumor grade, more specifically referred to as the “grade group”, which is determined by microscopic histopathologic examination (4, 5). While patients diagnosed with low grade prostate tumors have a 99% 5-year survival rate, patients with higher grade tumors and those who present with distant metastasis have significantly decreased survival (6). Standard treatment for prostate cancer includes active surveillance for patients with low grade tumors, localized therapy (radical prostatectomy and/or radiation) for intermediate and selected high grade tumors, and hormone therapy for patients with recurrence or metastatic disease (3). Despite the prognostic correlation with tumor grade group, racial disparities further contribute to prostate cancer patient outcomes. For example, Black males having a poorer prognosis compared to White males even when diagnosed with low grade prostate tumors (7–10). One critical knowledge gap in prostate cancer biology is the molecular events underlining higher incidence and mortality rates within this patient population, which could lead to better predictive biomarkers and personalized therapy.

N-linked glycosylation is a co-translational event necessary for cell surface, secreted, and circulating proteins (11, 12), wherein glycoconjugates containing N-acetylglucosamine (GlcNAc) are covalently attached to asparagine residues on the nascent carrier protein, followed by sequential addition of monosaccharides such as mannose, fucose, sialic acid, or GlcNAc (13, 14). Several biological processes are regulated by N-linked glycosylation including cell adhesion, immune modulation, cell-matrix interactions, and cell proliferation (15–19). Recent glycomic and proteomic studies have revealed extensive alterations in both the N-glycan profile and glycosyltransferase expression of several human cancers, including breast, lung, and prostate (20–22). Moreover, aberrant N-glycosylation has been shown to directly facilitate epithelial-to-mesenchymal transition (EMT) and subsequent metastatic protentional of cancer cells by directly altering the activity of extracellular matrix proteins and growth factor signaling (23). Given the role of N-glycosylation during EMT and metastasis, defining the N-glycome of prostate tumors can provide insight into the molecular mechanisms driving prostate cancer progression and can be used to discover new biomarkers or potential novel therapies.

Matrix-assisted laser desorption/ionization-mass spectrometry imaging (MALDI-MSI) is a new and innovative technique in glycobiology that can be used to profile N-glycans with spatial distribution in formalin-fixed paraffin-embedded (FFPE) samples and high throughput analysis of tissue microarrays (TMA) (24–26). This novel approach uniquely utilizes 1) the enzyme peptide-N-glycosidase F (PNGase F) that cleaves bound N-linked glycans from glycoproteins *in situ*, and 2) α-cyano-4-hydroxycinnamic acid (CHCA) ionization matrix for detection of N-linked glycans by MALDI-MSI (27). Previous studies have revealed distinct alterations in the N-glycan distribution between normal and prostate tumor tissue (22, 24, 28), with several of the N-glycan species elevated in prostate cancer being linked to EMT and metastasis (29–32). Current MALDI-MSI analyses of prostate cancer tissues have utilized large prostate tissue sections, and elegantly describe the N-glycan spatial differences between tumor and nontumor regions (22, 24, 28). Given these recent findings and the role of N-glycans in EMT, we hypothesized that N-glycan profiling may have potential to both define tumor grade and predict overall patient outcome in prostate cancer. We performed MALDI-MSI analysis on FFPE prostate cancer TMAs constructed from archived human prostate tissues from over 100 patients treated at the Markey Cancer Center. This patient data set included both cancer and matched normal tissue from racially and geographically diverse patients with over 10 years of follow-up data, allowing us to evaluate N-glycans as prognostic indicators for the clinical course of prostate cancer progression.

We observed significant N-glycan dysregulation between benign prostate tissue and tumor prostate tissue with several glycans tracking either positively or negatively with tumor grade group. Specifically, high mannose as well as tri- and tetra-antennary glycans were more abundant within tumor tissue and correlated with increasing tumor grade. Further, we expanded our analyses to access glycosylation patterns in populations disproportionally affected by prostate cancer. We report that glycosylation does not differ between patients from Appalachian and non-Appalachian populations in Kentucky; however, we found striking differences in the N-glycan profiles of low stage prostate cancer tumors between Black and White patients. Moreover, these glycans also predict opposing survival outcomes between Black and White patients. This surprising data highlights fundamental differences in carbohydrate metabolism during early tumorigenesis between the Black and White patient populations. Overall, our data suggest that aberrant N-linked glycosylation contributes to prostate cancer progression, which highlights the clinical potential of MALDI-MSI analysis for novel biomarker discover, and emphasizes the need for personalized medicine for prostate cancer patients.

## Results

### Utilizing TMAs for high-throughput analysis of prostate cancer N-glycans by MALDI-MSI

Previous MALDI-MSI analyses of prostate cancer tissue sections have revealed distinct differences in the spatial distribution of several species of N-glycans between tumor and nontumor regions (24, 28). We aimed to expand on these observations and utilize MALDI-MSI analysis to define the N-glycome of over 100 prostate cancer patients with demographical information and clinical course. We obtained FFPE prostate TMAs containing both benign prostate tissue and prostate tumor tissue with over 10 years of patient follow-up data from the Biospecimen Procurement and Translation Pathology Shared Resource Facility (BPTP SRF) of the Markey Cancer Center (Lexington, KY). The TMAs analyzed included patient samples of prostate cancer grade groups 1 through 5, and clinicopathological parameters included race, geographic location, as well as disease recurrence and patient survival (**Supplemental Table 1**). Utilizing the modern grading system, few patients are diagnosed with grade group 4 prostate cancer at radical prostatectomy (33), and this fact is reflected in our cohort of patients with only three grade group 4 patient samples. Therefore, these samples were omitted from the analysis due to a low statistical power. TMA slides were prepared using previously established MALDI-MSI workflow (24, 27). First, bound N-glycans were cleaved from glycoproteins by the addition of PNGaseF; then, CHCA, an ionization matrix was applied uniformly using the HTX high velocity dry-spraying robot (34). Released N-glycans were analyzed using a Waters Synapt G2 ion-mobility enabled mass spectrometer equipped with an Nd:YAG UV laser (**Figure 1A**). Ion mobility improved glycan detection by separating N-glycans from ionization matrix based on differential collision cross section (**Figure 1B**). Using this method, we detected 46 N-glycans across all tissue samples (**Figure 1C** and **Supplemental Table 2**). Representative HDI images of the four most abundant N-glycans (1501, 1663, 1809, and 1976 m/z) are shown in **Figure 1D-G**. Interestingly, these biantennary complex N-glycans, while sharing a common core structure, show a wide range of abundance across the TMA.

**Figure 1.**
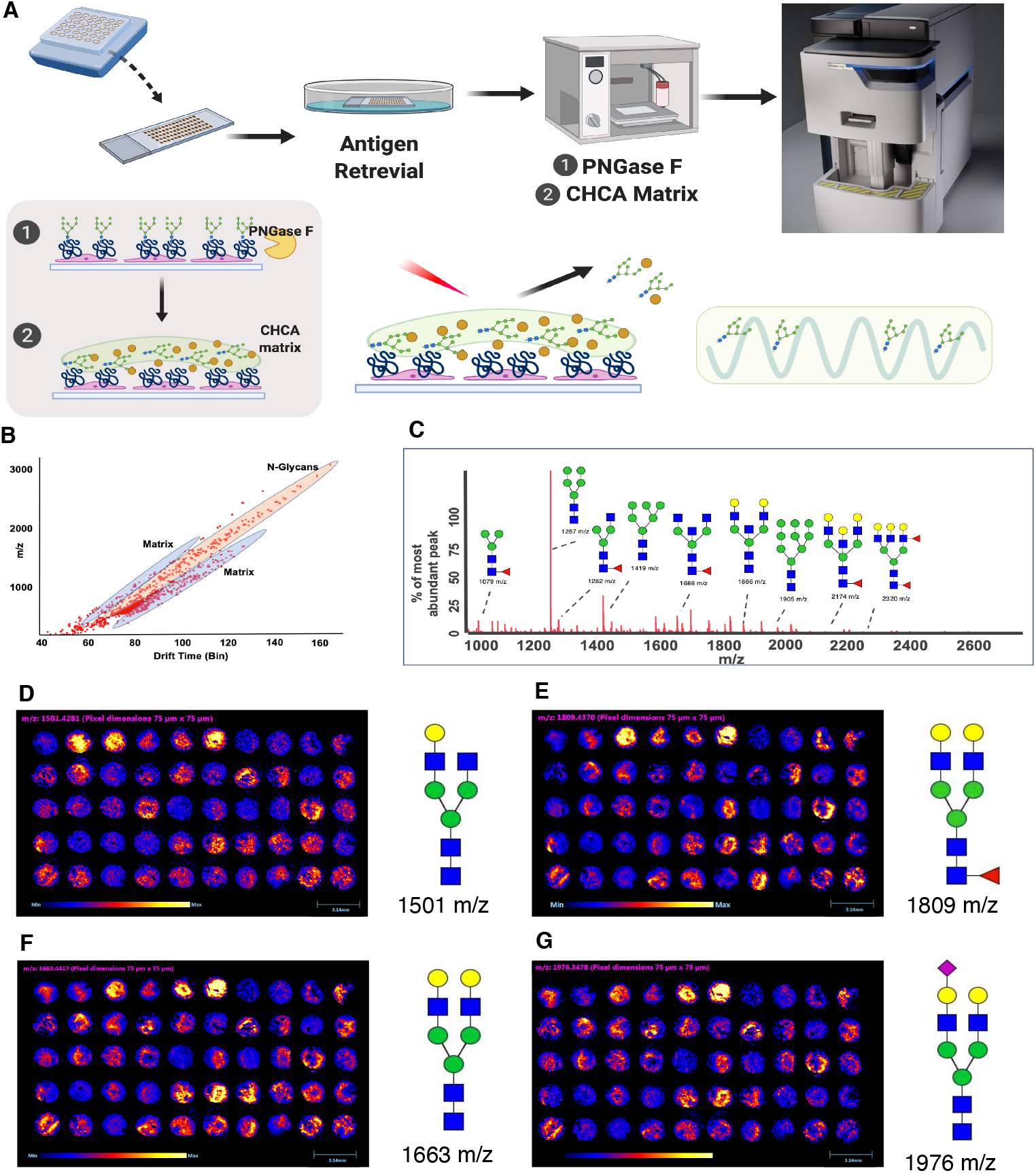
Overview of MALDI-MSI workflow for N-glycan detection in prostate cancer TMAs. (**A**) Schematic workflow of N-glycan imaging using MALDI coupled to high resolution mass spectrometry. In brief, FFPE TMAs were sectioned to 5μm followed by dewaxing and antigen retrieval. Slides were treated with PNGase F to release N-glycans followed by CHCA (ionization matrix) application using an HTX high velocity sprayer. Glycans and matrix were ionized using a Nd:YAG UV laser and were separated by ion mobility. Glycan mass spectra were acquired by Waters Synapt G2 high resolution mass spectrometer. (**B**) Scatter plot of monoisotopic mass versus drift time in the ion mobility cell for N-glycans and the MALDI matrix. (**C**) Extracted ion chromatogram of released N-glycans based on ion mobility separation. Structures of several detected glycans are shown on the plot. (**D**) HDI images of 1501 m/z, (**E**) 1809 m/z, (**F**) 1663 m/z, and (**G**) 1976 m/z. Intensity and size scales are located beneath the image. Blue: least abundant, Yellow: most abundant. Glycan structures are placed on the right side of their corresponding image. Structure key: blue square-N-acetylglucosamine, green circle-mannose, yellow circle-galactose, purple diamond-sialic acid, and red triangle-fucose. Scale bar-3.14mm.

### Core fucosylated, bisecting, and sialylated N-glycans are predominantly found in benign tissue

Core fucosylation is an important N-glycan modification wherein a fucose moiety is added via α1,6-linkage to the innermost GlcNAc residue, altering the activity of the attached protein. Specifically, fucosylation of cell membrane receptors and proteins such as EGFR, TGFβ, E-cadherin, and integrins influences ligand binding, receptor dimerization, and signaling capacities (35–38). Core fucosylation is frequently increased in tumor tissue and inhibiting the addition of a core fucose moiety to glycans reduces cancer progression (39). Despite these findings, we observed a decrease in both the core fucosylated (1485, 1647, and 1809 m/z) (**Figure 2A-C**) and non-fucosylated (1339, 1501, and 1663 m/z) (**Figure 2D-F**) structures of several biantennary N-glycans in prostate tumor tissue compared to benign. Further, the abundance of these specific N-glycans decreased with tumor grade group. These results suggest that reductions in certain core fucosylated glycan species are part of N-glycan reprogramming in prostate cancer and may play a role in prostate cancer tumorigenesis.

**Figure 2.**
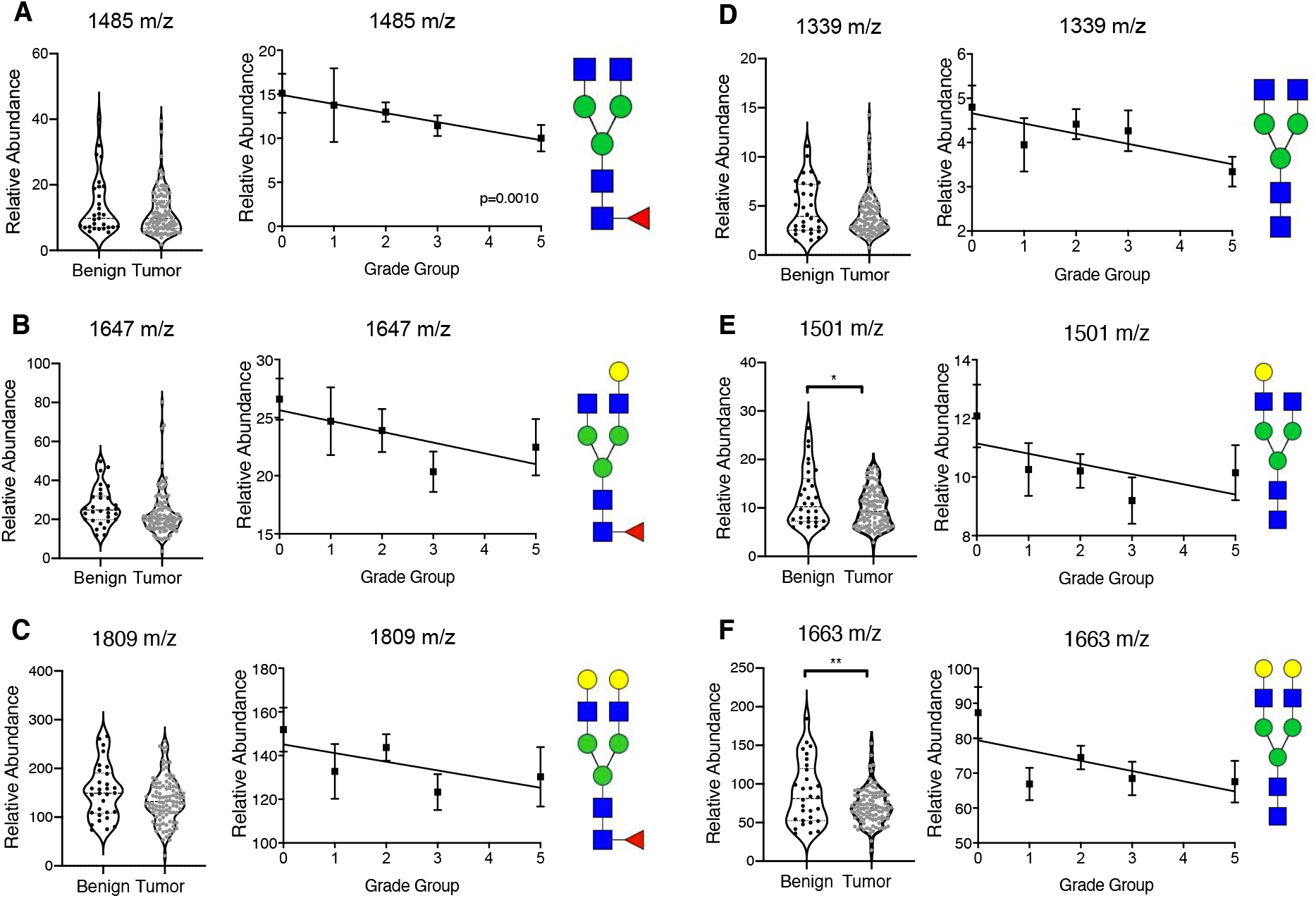
Prostate tumor tissue exhibits decreased abundance of biantennary N-glycans with and without a core fucose. N-glycan relative abundance for benign versus grouped prostate tumor tissue (left), relative abundance stratified by tumor grade (middle), and representative structure (right) for biantennary N-glycans with: (**A**) 1485 m/z, (**B**) 1809 m/z, (**C**) 1647 m/z and with core fucose modification: (**D**) 1339 m/z, (**E**), 1501 m/z, and (**F**) 1663 m/z. Values represent mean ± S.E.M. analyzed by student’s t-test (benign vs grouped tumor tissue) or simple linear regression. Benign (n=30), grade group 1 tumors (n=21), grade group 2 tumors (n=48), grade group 3 tumors (n=21), and group grade 5 tumors (n=15). p<0.05. Structure key: blue square-N-acetylglucosamine, green circle-mannose, yellow circle-galactose, purple diamond-sialic acid, and red triangle-fucose.

Bisecting N-glycans are another class of N-glycans that play a role in cancer progression. This class is characterized by the addition of a β1,4-linked GlcNAc to the β-mannose residue of the glycan core, which inhibits further processing and elongation by glycosyltransferases (26, 40, 41). Accumulation of bisecting N-glycans correlates to anti-tumorigenic potential in cancer cells by limiting the formation of larger complex glycan structures (42, 43). Consistent with this observation, we found that two bisecting N-glycans (1542 and 1704 m/z) predominantly accumulated in benign prostate tissue compared to prostate tumor tissue, which decreased with tumor grade group (**Supplemental Figure 1A-B**); one bisecting N-glycan remained unaffected (1866 m/z) (**Supplemental Figure 1C**). Our findings support the notion that bisecting N-glycans are anti-tumorigenic and play a protective role in cancer progression.

Sialic acids are terminal monosaccharides on N-glycans that project into the extracellular environment, and play an important role in protein-protein interaction and cellular recognition (44). Sialylation is one of the building blocks to form the cancer-associated sialyl-Lewis antigens, which have been shown to be involved in cell adhesion by selectin interaction (45, 46). Moreover, sialylation in tumor tissue is known to be a mechanism of resistance to cell death and may protect cells from infiltrating immune cells, thereby inhibiting immune surveillance (47). We found that two sialylated N-glycans (1976 and 2122 m/z) were more abundant in benign prostate tissue compared to prostate tumor tissue, which decreased with tumor grade group (**Supplemental Figure 1D-E**); one sialylated N-glycan was unchanged (2341m/z) (**Supplemental Figure 1F**). These data suggest N-glycan sialylation is not a major driver of immune surveillance or modulating the microenvironment immune landscape during prostate cancer progression.

### Prostate tumors exhibit increased high-mannose and branched complex N-glycans in a grade group-dependent manner

Previous MALDI-MSI analyses of prostate cancer tissue sections have revealed distinct differences in the spatial distribution of several species of N-glycans between tumor and nontumor regions (24, 28). Specifically, high-mannose glycans were almost exclusively detected in prostate tumor tissue. We aimed to expand on these observations and identify glycans that accumulate with increased tumor grade group. Consistent with previous findings, prostate tumor tissue exhibited higher abundance of several high-mannose N-glycans (1581, 1743, and 1905 m/z) compared to benign prostate tissue, which increased with tumor grade group (**Figure 3A-C**). High-mannose glycans are produced early in the N-glycan biosynthetic pathway, wherein mannose residues are sequentially added to the growing glycan chain in the ER, followed by further processing by mannosidases and glycotransferases into more structurally diverse complex and hybrid glycans in the Golgi apparatus (48, 49). High-mannose glycans are routinely detected in high abundance in cancer tissues and have been implicated in the progression of several human cancers including liver, lung, and breast (50–52). Together, combined with previous studies, our data highlights the clinical relevance of high-mannose glycans in prostate cancer. Additionally, these data suggest prostate tumors exhibit either incomplete N-glycan biosynthesis or enhanced mannose metabolism that contributes to prostate cancer progression.

**Figure 3.**
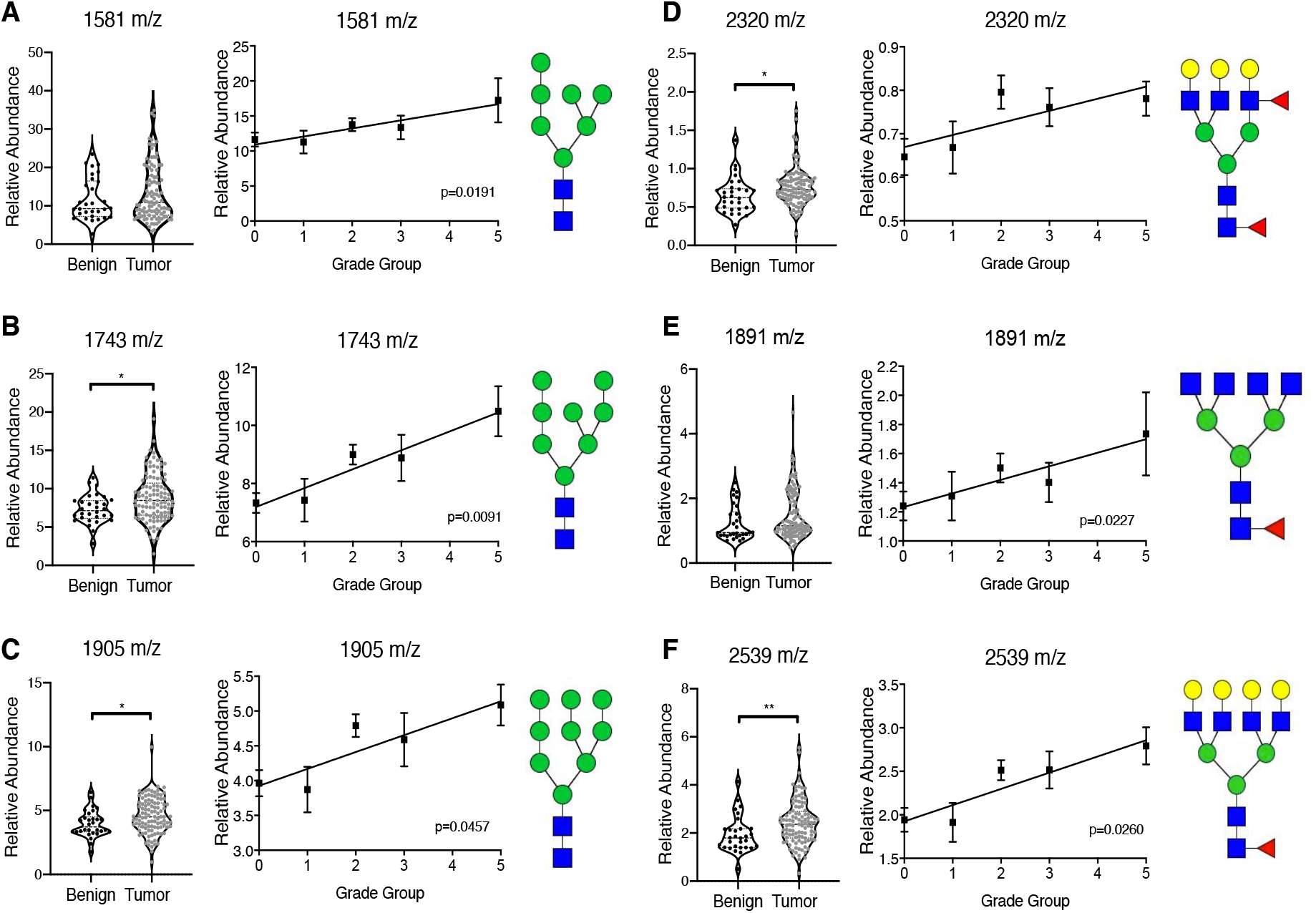
Prostate tumor tissue exhibits increased abundance of high mannose as well as tri- and tetra-antennary N-glycans proportional to tumor grade. N-glycan relative abundance for benign versus grouped prostate tumor tissue (left), relative abundance stratified by tumor grade group (middle), and representative structure (right) for high mannose N-glycans: (**A**) 1581 m/z, (**B**) 1743 m/z, (**C**) 1905 m/z and complex tri-/tetra-antennary N-glycans: (**D**) 2320 m/z, (**E**), 1891 m/z, and (**F**) 2539 m/z. Benign (n=30), grade group 1 tumors (n=21), grade group 2 tumors (n=48), grade group 3 tumors (n=21), and grade group 5 tumors (n=15). Values represent mean ± S.E.M. analyzed by student’s t-test (benign vs grouped tumor tissue) or simple linear regression. p<0.05 and **p<0.01. Structure key: blue square-N-acetylglucosamine, green circle-mannose, yellow circle-galactose, purple diamond-sialic acid, and red triangle-fucose.

Other glycans that correlated positively with tumor grade group include several branched complex N-glycans (2320, 1891, and 2539 m/z) with a core fucose residue (**Figure 3D-F**). Increased tri- and tetra-antennary branched glycan structures have been linked to many aspects of tumorigenesis including neoplastic transformation, cell proliferation, and abnormal cell morphology (26, 53). Moreover, it’s been demonstrated that the addition of tri- and tetra-antennary branched glycans to E-cadherin impairs cell adhesion and promotes tumor cell invasion (54). Our findings suggest increased prostate cancer tumorigenesis is through similar mechanisms. Notably, core fucosylated biantennary complex glycans are lower in prostate tumor tissue and decrease with tumor grade group (**Figure 2A-C**), while tri- and tetra-antennary complex glycans show an opposing trend (**Figure 3D-F**). These data suggest glycan branching, rather than core fucosylation, plays a greater role in prostate cancer tumorigenesis.

### Elevated high mannose and complex N-glycans in prostate tumor tissue are not prognostic markers for disease progression across all patient populations

Higher tumor grade groups are typically associated with poorer patient outcomes in prostate cancer (4, 6). Therefore, we hypothesized that high mannose and branched complex N-glycans would predict the clinical course of prostate cancer progression. To assess this, we took advantage of the 10 year follow-up data linked to the patient samples on the prostate TMA. First, we analyzed the relative abundance of each glycan with respect to whether or not that patient had disease recurrence. Surprisingly, we found that the relative abundance of both species of N-glycans was not altered between patients who did not have disease recurrence compared to those with local or regional recurrence (**Figure 4-F**). Second, we assessed the relative abundance of each glycan with overall survival for each patient. Linear regression analysis revealed that increased abundance of high-mannose and branched complex N-glycans did not correlate to better or poorer overall survival for patients (**Figure 4G-L**). To our surprise, even though high mannose and branched complex glycans correlated positively with tumor grade group, they did not correlate with recurrence and overall survival across all patient populations. These data suggest that there are other co-founding factors that contribute to prostate cancer progression. It is well known that health disparities exist within prostate cancer patient cohorts, specifically in men from rural Appalachia and Black men (7–10, 55–57). Thus, we hypothesize that these distinct patient populations have unique glycan signatures that contribute to the lack of correlation between glycan abundance and patient outcome in our initial analysis.

**Figure 4.**
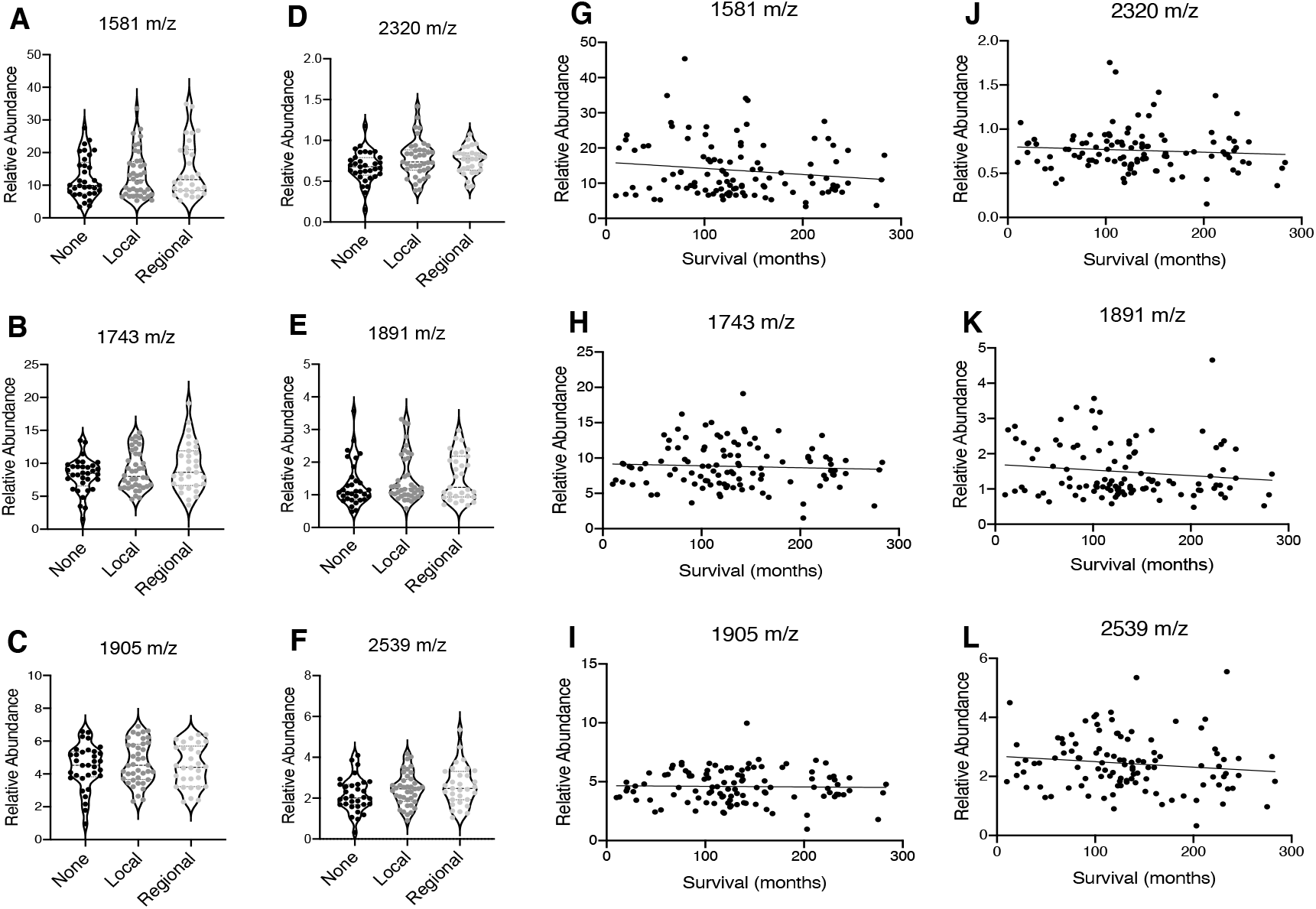
Elevated high mannose and complex N-glycans in prostate tumor tissue are not prognostic markers for disease progression across all patient populations. N-glycan relative abundance stratified by disease recurrence for high mannose N-glycans: (**A**) 1581 m/z, (**B**) 1743 m/z, (**C**) 1905 m/z, and complex tri-/tetra-antennary N-glycans: (**D**) 2320 m/z, (**E**), 1891 m/z, and (**F**) 2539 m/z. Data are plotted as individual patient samples, analyzed by One-way ANOVA. No recurrence/None (n=33), local recurrence (n=40), and regional recurrence (n=31). Correlation of relative abundance to overall patient survival for high mannose N-glycans: (**G**) 1581 m/z, (**H**) 1743 m/z, (**I**) 1905 m/z, and complex tri-/tetra-antennary N-glycans: (**J**) 2320 m/z, (**K**), 1891 m/z, and (**L**) 2539 m/z. Data are plotted as individual patient samples, analyzed by simple linear regression analysis (n=108).

### N-glycosylation does not contribute to poorer survival in prostate cancer patients from rural Appalachia

Several epidemiological studies have revealed prostate cancer patients from rural Appalachia have poorer overall survival despite having lower incidence compared to patients from non-Appalachian counties (56). Yet, the molecular mechanisms underlining this health disparity are largely unknown. As our patient cohort includes patients within Kentucky residing in Appalachian and non-Appalachian counties, we hypothesized N-glycan dysregulation contributes to poorer outcomes for patients from rural Appalachia. We compared the relative abundance of all 46 N-glycans between Appalachian and non-Appalachian patients by grade group, and found no significant differences in N-glycan abundance for several species of N-glycans, including high-mannose (**Supplemental Figure 2A**), bisecting (**Supplemental Figure 2B**), sialylated (**Supplemental Figure 2C**), or core fucosylated (**Supplemental Figure 2D**). Abundance of these specific glycans is not altered between patients who did not have disease recurrence compared to those with local or regional recurrence for Appalachian status (**Supplemental Figure 3A-D**). Moreover, we did not observe any significant difference in the correlation between glycan abundance and overall survival between Appalachian versus non-Appalachian patients (**Supplemental Figure 3A-D**). Our data suggest N-glycosylation does not contribute to prostate cancer progression in Appalachian patients, and it supports the notion that late diagnosis, rather than underlining molecular features, drives the health disparity between Appalachian and non-Appalachian prostate cancer patients (56).

### N-glycosylation contributes to the health disparity in Black prostate cancer patients

While increasing evidence suggests that molecular and genetic alterations contribute to the racial disparity between Black and White prostate cancer patients (58–61), how specific differences in tumor biology drive accelerated disease progression in Black men remains largely unknown. Our TMAs included a cohort of black patients treated at the Markey Cancer Center. Therefore, we expanded our analysis to examine the N-glycan profile of White and Black prostate cancer patient samples, and whether those differences contributed to the health disparity in Black men. We analyzed the relative abundance of all 46 N-glycans detected between Black and White patient samples, stratified by grade group, and found 9 structurally diverse glycans that were elevated in grade group 1 tumors of Black patients. These N-glycans included pauci-mannose (**Figure 5A, B**), biantennary complex (**Figure 5C**), sialylated (**Figure 5D**), core fucosylated (**Figure 5E, F**), bisecting (**Figure 5G, H**), and tetra-antennary complex (**Figure 5I**) N-glycans. Several of the glycans elevated in low grade tumors from Black patients compared to White were unchanged in benign tissue. Notably, high-mannose and complex glycans that were found to be elevated in prostate tumor tissue from our initial analysis (**Figure 3**) were not significantly altered between Black and White patient samples (**Supplemental Figure 4A-E**). This finding suggests we have identified a glycan signature that is unique to low grade tumors from Black patients, which could inform novel early diagnostic strategies and novel treatment strategies for this patient population. We stratified relative abundance for all nine glycans elevated in grade group 1 tumors by recurrence between Black and White patients, and did not observe any significant difference in glycan abundance between Black and White patients who had local disease recurrence compared to those with none (**Supplemental Figure 5**). Our patient cohort did not include tumor samples from Black patients with regional recurrence, thus we excluded regional recurrence form our analyses between Black and White patients. Strikingly, six of the nine glycans that are accumulated in low grade tumors predict opposing overall survival between Black and White patients (**Figure 6**). This surprising data suggests the fundamental changes in N-glycan metabolism that occur during early tumorigenesis between the White and Black patients potentially contribute to the health disparity in prostate cancer disease progression.

**Figure 5.**
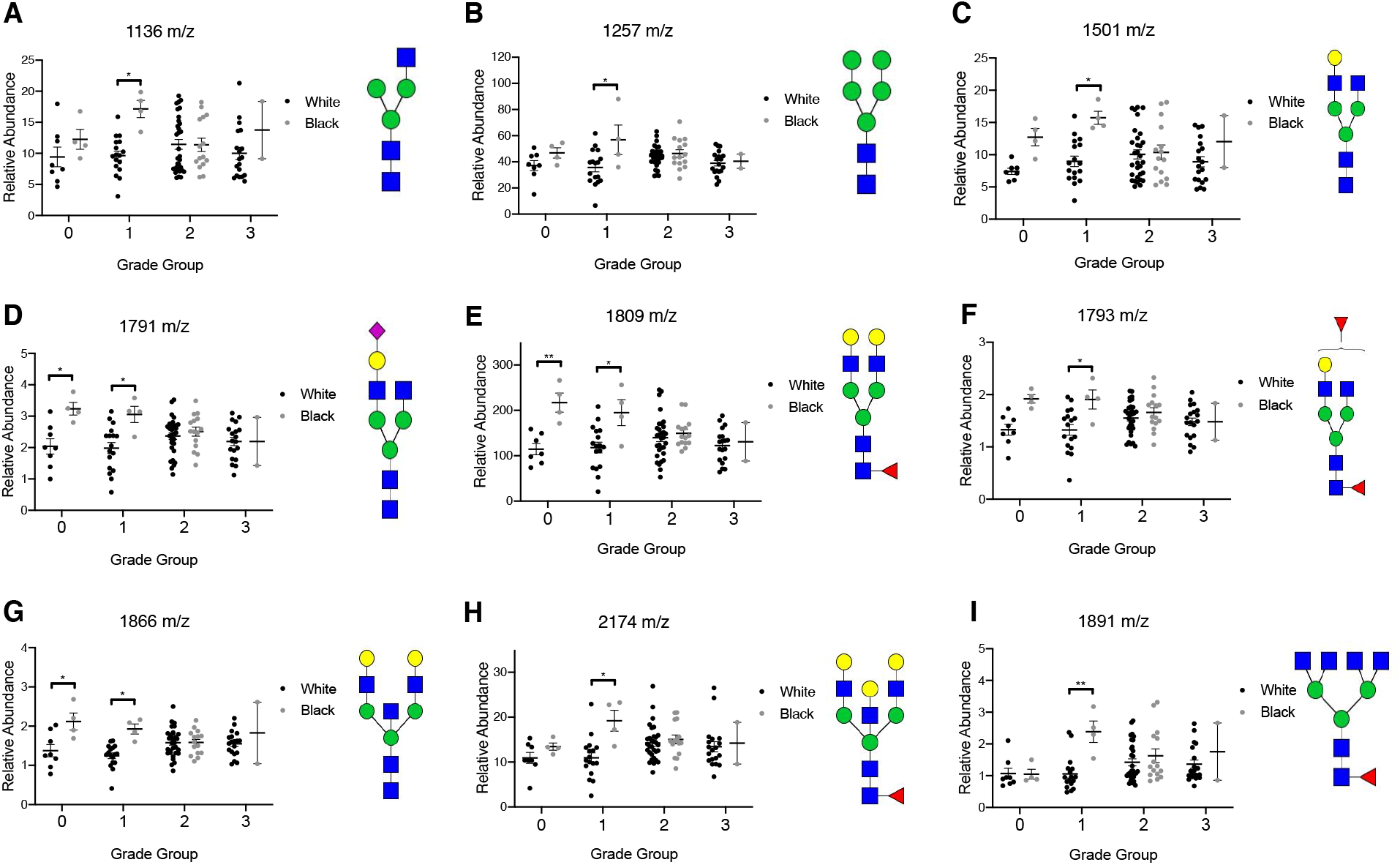
Low grade prostate cancer tumors between Black and White patients exhibit significantly different N-glycan profiles. N-glycan relative abundance for White versus Black patient samples stratified by tumor grade (left), and representative structure (right) for (**A**) 1136 m/z (pauci-mannose), (**B**) 1257 m/z (pauci-mannose), (**C**) 1501 m/z (bi-antennary complex), (**D**) 1791 m/z (sialylated), (**E**) 1809 m/z (core fucosylated), (**F**) 1793 m/z (core fucosylated), (**G**) 1866 m/z (bisecting), (**H**) 2174 m/z (bisecting/core fucosylated), and (**I**) 1891 m/z (tetra-antennary complex/core fucosylated). Error bars represent mean ± S.E.M. analyzed by Two-way ANOVA. White: benign (n=8), grade group 1 tumors (n=17), grade group 2 tumors (n=31), and grade group 3 tumors (n=19). Black: benign (n=4), grade group 1 tumors (n=4), grade group 2 tumors (n=15), and grade group 3 tumors (n=2). p<0.05 and **<0.01. Structure key: blue square-N-acetylglucosamine, green circle-mannose, yellow circle-galactose, purple diamond-sialic acid, and red triangle-fucose.

**Figure 6.**
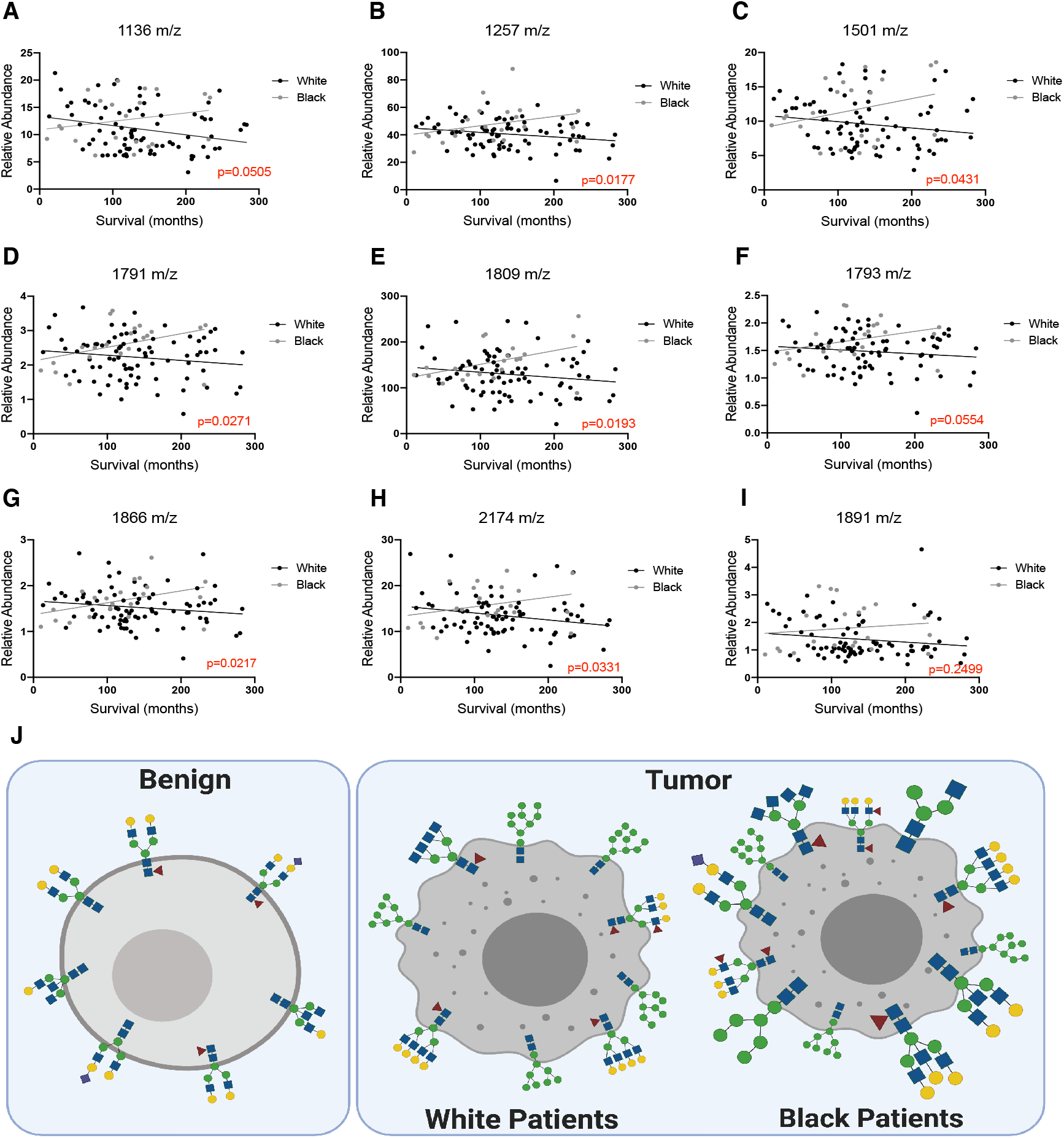
N-glycan status of low grade prostate tumors of White and Black patients predict opposing trends in overall survival. Correlation of relative abundance to overall patient survival for (**A**) 1136 m/z (pauci-mannose), (**B**) 1257 m/z (pauci-mannose), (**C**) 1501 m/z (bi-antennary complex), (**D**) 1791 m/z (sialylated), (**E**) 1809 m/z (core fucosylated), (**F**) 1793 m/z (core fucosylated), (**G**) 1866 m/z (bisecting), (**H**) 2174 m/z (bisecting/core fucosylated), and (**I**) 1891 m/z (tetra-antennary complex/core fucosylated). Data are plotted as individual patient samples, analyzed by simple linear regression analysis. White (n=81) and Black (n=21). p values are displayed in red for each glycan. (**J**) Illustration of the N-glycan profile of benign prostate tissue compared to low grade tumors from White and Black prostate cancer patients.

## Discussion

An increasing body of evidence suggests that aberrant N-glycosylation plays a key role in several aspects of tumorigenesis, such as tumor cell invasion and metastasis, cell-matrix interactions, tumor angiogenesis, and cell signaling and communication (20). With the advent of new high throughput mass spectrometry based technologies, such as MALDI-MSI, N-linked glycomic profiling of patient tumor tissues has demonstrated remarkable potential for early diagnosis, risk prediction, and treatment outcome for several cancers (62). Moreover, MALDI-MSI analysis of N-glycans provides insight into the function of N-linked glycosylation in tumor metabolism and cancer progression (63, 64). The use of TMAs is advantageous for high-throughput investigation during a single experiment using widely available FFPE patient samples, often including clinical follow-up data. In the present study, we utilized prostate cancer TMAs including benign and tumor tissue resected from over 100 patients with 10 years of clinical follow-up data to perform N-glycan profiling by MALDI-MSI analysis. We found specific glycan signatures between benign compared to prostate tumor tissue. Further, we have identified a unique glycan profile in low grade tumors from Black compared to White prostate cancer patients, which potentially contributes to the racial disparity in prostate cancer.

We observed significant dysregulations in multiple species of N-glycans between benign prostate tissue and prostate tumor tissue. Specifically, prostate tumors exhibit accumulation of high-mannose glycans which increase with tumor grade group (**Figure 3A-C**), a common feature of human cancers that correlates with more aggressive cancer phenotypes (20, 21). Accumulation of high mannose N-glycans in prostate tumors suggests a lack of N-glycan trimming reactions and a decrease in mannosidase activity, or aberrant mannose metabolism (50–52). Additionally, we found prostate tumors accumulate tri- and tetra-antennary complex glycans containing a core fucose moiety (**Figure 3D-F**), suggesting prostate tumors have enhanced GlcNAc metabolism. N-glycan β1,6-branching, which gives rise to these structures, has been implicated in in several tumorigenic processes including neoplastic transformation, cell proliferation, and abnormal cell morphology (26, 53). Our findings suggest that increased N-glycan β1,6-branching and the accumulation of high-mannose glycans contribute to prostate cancer progression. Despite high-mannose and branched N-glycans increasing with tumor grade group, these specific glycans were not able to predict the clinical course of prostate cancer progression over a 10 year follow-up interval in the present study (**Figure 4**). Future analyses should be extended to longer follow-up intervals and a wider spread of clinical behaviors (respons to therapy, co-morbidities, etc.) to define the prognostic value of these specific glycans in prostate cancer. We also found several species of N-glycans were elevated in benign tissue compared to prostate cancer tumors, including core fucosylated, bisecting, and sialylated glycans (**Figure 2, Supplemental Figure 1**). Strikingly, biantennary complex glycans with a core fucose moiety are lower in prostate tumor tissue and decrease with tumor grade group (**Figure 4A-C**), while tri- and tetra-antennary core fucosylated glycans are increased (**Figure 3D-F**), suggesting branching, rather than core fucosylation, contributes to prostate cancer progression.

Health disparities among different prostate cancer patient populations have been well documented, with men from rural Appalachia and Black men having higher mortality rates (7–10, 56). Yet, the molecular mechanisms driving poorer patient outcomes in these distinct populations are largely unknown. Our patient cohort is unique in that it includes samples from two disproportionally affected populations. We found N-glycan status is not a good classifier to separate the Appalachian and non-Appalachian patient population, which supports the hypothesis that late diagnosis, or other confounding factors, contribute to the health disparity in this population (**Supplemental Figure 2 and 3**). Conversely, we found several structurally diverse N-glycans that differentially accumulated between White and Black prostate cancer patients (**Figure 5**), that were distinct from the high-mannose and complex glycans that were found to be elevated in prostate tumor tissue from our initial analysis (**Figure 3**, **Supplemental Figure 4A-E**). Furthermore, these glycans predict opposing overall survival between each patient population (**Figure 6**). This finding suggests we have identified a glycan signature that is unique to low grade tumors from Black patients, which could be an important prognostic tool to predict stage 1 prostate tumors in the black population, hence warranting future validation. These data also raise the concern that Black and White patients would potentially respond to treatment options differently, and/or whether more targeted therapeutics are required for Black patients. Future experiments should include defining the molecular perturbations of N-glycan metabolism in Black prostate cancer patients, as identifying these features could lead to key to developing novel disease biomarkers and personalize therapies for this disproportionally affected population.

In summary, our data suggest that aberrant N-linked glycosylation contributes to prostate cancer progression, and identifies high-mannose, as well as branched glycans, as potential disease biomarkers. Moreover, our study is the first to define the N-glycan profiles between prostate tumors from Black and White patient populations, and suggest differences in N-glycosylation is a molecular feature in low grade tumors that potentially contributes to the health disparity in prostate cancer disease progression. Overall, these results warrant investigation to define glycan metabolism and the regulatory mechanisms that contribute to aberrant protein glycosylation in prostate cancer, with an emphasis on defining the unique features of low grade prostate tumors from Black patients, as they relate to patient prognosis. Future studies should be expanded to include glycoproteomic analysis to define the specific proteins that are differentially glycosylated. Such studies can provide insight into the molecular drivers of prostate cancer progression and health disparities, which can be used to discover new biomarkers and novel personalized therapies.

### Limitations of Study

This study employs cutting edge mass spectrometry imaging to identify tumor specific and patient demographic alterations in N-glycosylation in prostate cancer. While MALDI-MSI is a powerful tool for high-throughput N-glycan profiling of a large number of patient samples, we are still limited by the patient cohort selected for this study. For our targeted demographic analysis, sample size was small for several groups, thus future studies should include more patients to confirm our findings. Moreover, we analyzed prostate tumors from small tissue cores rather than larger tissue sections containing both tumor and nontumor stroma regions. As many tumor cores are not purly tumor tissue, this could contribute to increase variance in our results. future analyses should be expanded to define N-glycosylation in different tumor regions defined by microenvirental pressure in larger prostate tumor tissue sections.

## Methods

### Chemicals and Reagents

High-performance liquid chromatography-grade acetonitrile, ethanol, methanol, water, alpha-cyano-4-hydroxycinnamic acid (CHCA) and Trifluoroacetic acid (TFA) were purchased from Sigma-Aldrich. Histological-grade xylenes was purchased from Spectrum Chemical. Citraconic anhydride for antigen retrieval was obtained from Thermo Fisher Scientific. Recombinant PNGaseF Prime was obtained from Bulldog Bio, Inc. (Portsmith, NH, USA).

### Clinical Prostate Cancer FFPE Tissue Microarrays

Tissue microarrays (TMAs) were created from residulal FFPE radical prostatectomy samples by the Biospecimen Procurement and Translation Pathology Shared Resource Facility (BPTP SRF) of the Markey Cancer Center (MCC) with approval from the institutional review board (IRB). These specimens were coupled with de-identified demographic and clinical data provided by the Cancer Research Informatics (CRI) SRF and the MCC with approval from the IRB. The TMAs contained prostate tumor tissue (n=112 samples) and benign prostate tissue (n=30 samples) from 112 patients. 21/112 patient tumor samples were grade group 1; 48/112 were grade group 2; 21/112 were grade group 3; 3/112 were grade group 4; and 15/112 were grade group 5. **Supplemental Table 1**. All tissues were de-identified to the analytical investigators.

### Tissue Preparation and Enzyme Digestion

FFPE TMA slides were processed as previously described (1, 2). In brief, tissues were dewaxed and rehydrated followed by antigen retrieval in citraconic anhydride buffer (25μl citraconic anhydride, 2μl 12 M HCl, and 50μl HPLC-grade water, pH 3.0-3.5). Recombinant PNGase F (0.1μg/μl) was applied using an M5 TMSprayer Robot (HTX Technologies LLC, Chapel Hill, NC). Enzyme was sprayed onto the slide at a rate of 25μl /min with a 0mm offset and a velocity of 1200mm/min at 45°C and 10psi for 15 passes, followed by a 2-hour incubation at 37°C in a prewarmed humidity chamber.

After incubation, slides were desiccated and 7mg/ml CHCA matrix in 50% acetonitrile with 0.1% TFA was applied at 0.1ml/min with a 2.5mm offset and a velocity of 1300mm/min at 79°C and 10psi for 10 passes using the M5 Sprayer. Slides were stored in a desiccator or immediately used for MALDI-MSI analysis.

### N-Glycan MALDI-MSI Analysis

A Waters Synapt G2Si mass spectrometer (Waters Corporation, Milford, MA) equipped with an Nd:YAG UV laser with a spot size of 50um was used to detect released N-glycans at X and Y coordinates of 75um. Data acquisition, spectrums were uploaded to High Definition Imaging (HDI) Software (Waters Corporation) for mass range analysis from 750 to 3500m/z. For N-glycan quantification, regions of interest (ROI) were defined for each patient sample on the TMAs using HDI image ROI drawing tool. For all pixels defined within a ROI, peak intensities were averaged and normalized by total ion current. HDI generated glycan images were obtained for most abundant N-glycans detected across all patient samples. Representative glycan structures were generated in GlycoWorkbench.

### Statistics

Statistical analyses were carried out using GraphPad Prism. All numerical data are presented as individual data points or mean ± S.E.M. Grouped analysis was performed using two-way ANOVA. Column analysis was performed using one-way ANOVA or unpaired t-test when appropriate. XY analysis was performed using simple linear regression. A p-value less than 0.05 was considered statistically significant.

### Study Approval

TMAs containing human prostate tissue were created by the BPTP SRF of the MCC with approval from the IRB. Samples were coupled with de-identified demographic and clinical data provided by the CRI SRF of the MCC with approval from the IRB. Use of the tissue and de-identified information for the purpose of this study was given an exempt status from the IRB.

## Author Contributions

RCS and DBA conceptualized the study and designed experiments. LRC, LEAY and AES performed the experiments. LRC, JL, MSG, and GLA analyzed the data and generated figures. LRC, GLA, MSG, and RCS wrote the manuscript. All authors read and approved the manuscript.

## Acknowledgements

This study was supported by NIH grant R01 AG066653, St Baldrick’s Career Development Award, and Rally Foundation Independent Investigator Grant. This research was also supported by funding from the University of Kentucky Markey Cancer Center and the NIH-funded Biospecimen Procurement & Translational Pathology Shared Resource Facility, as well as the Cancer Research Informatics Shared Resource Facility, of the University of Kentucky Markey Cancer Center P30CA177558.

